# The Parkinson’s Phenome: Traits Associated with Parkinson’s Disease in a Large and Deeply Phenotyped Cohort

**DOI:** 10.1101/270934

**Authors:** Karl Heilbron, Alastair J. Noyce, Pierre Fontanillas, Babak Alipanahi, Mike A. Nalls, Paul Cannon

**Affiliations:** 23andMe, Inc., 899 W Evelyn Avenue, Mountain View, California 94041 USA; Preventive Neurology Unit, Wolfson Institute of Preventive Medicine, Queen Mary University of London, London, UK; Department of Molecular Neuroscience, UCL Institute of Neurology, London, UK; Data Tecnica International, Glen Echo, MD, USA; Laboratory of Neurogenetics, National Institute on Aging, Bethesda, USA

## Abstract

**Background:** Observational studies have begun to characterize the wide spectrum of phenotypes associated with Parkinson’s disease (PD), but recruiting large numbers of PD cases and assaying a diversity of phenotypes has often been difficult. Here, we set out to systematically describe the PD phenome using a cross-sectional case-control design in a large database.

**Methods:** We analyzed the association between PD and 840 phenotypes derived from online surveys. For each phenotype, we ran a logistic regression using an average of 5,141 PD cases and 65,459 age- and sex-matched controls. We selected uncorrelated phenotypes, determined statistical significance after correcting for multiple testing, and systematically assessed the novelty of each significant association. We tested whether significant phenotypes were also associated with disease duration in PD cases.

**Findings:** PD diagnosis was associated with 149 independent phenotypes. We replicated 32 known associations and discovered 49 associations that have not previously been reported. We found that migraine, obsessive-compulsive disorder, seasonal allergies, and anemia were associated with PD, but were not significantly associated with PD duration, and tend to occur decades before the average age of diagnosis for PD. Further work is needed to determine whether these phenotypes are PD risk factors or whether they share common disease mechanisms.

**Interpretation:** We used a systematic approach in a single large dataset to assess the spectrum of traits that were associated with PD. Some of these traits may be risk factors for PD, features of the pre-diagnostic phase of disease, or manifestations of PD pathology. The model outputs from all 840 logistic regressions are available to the research community and may be used to generate hypotheses regarding PD etiology.

**Funding:** The Michael J. Fox Foundation, Parkinson’s UK, Barts Charity, National Institute on Aging, and 23andMe, Inc.

**Research in Context:** *Evidence before this study:* We used PubMed to perform a MEDLINE database search for review articles published up to January 21^st^, 2018 that contained the keywords “Parkinson” and “epidemiology” in the title or abstract. We performed additional MEDLINE searches for each phenotype that was significantly associated with PD. Although dozens of phenotypes have been tested for an association with PD, only a few associations have been consistently repeatable (*e.g.* pesticide exposure, coffee consumption).

*Added value of this study:* We systematically tested for an association between PD and 840 phenotypes using up to 13,546 cases and 1·3 million controls, making this one of the largest PD epidemiology studies ever conducted. We discovered 49 novel associations that will need to be replicated or validated. We found 44 associations for phenotypes that have previously been studied in relation to PD, but for which an association has not been consistently demonstrated.

*Implications of all the available evidence:* Taken together with results from previous studies, this series of case-control analyses adds evidence for associations between PD and many phenotypes that are not currently thought to be part of the canonical PD phenome. This work paves the way for future studies to assess whether any of these phenotypes represent PD risk factors and whether any of these risk factors are modifiable.

## Introduction

The clinical diagnosis of PD is based on the canonical motor symptoms of bradykinesia, plus tremor and/or rigidity, but there has recently been a growing understanding of the importance of the non-motor aspects of PD.^1^ Epidemiological studies have uncovered a collection of traits, environmental exposures, and comorbidities (we will collectively refer to these as “phenotypes”) that are associated with PD, including reduced coffee consumption, reduced tobacco use, increased melanoma skin cancer, and increased serum uric acid levels.^2^ However, the prevalence of PD is approximately 1% in individuals over 65 years old^3^, making it difficult to amass the large numbers of cases required to detect more subtle phenotypic associations using traditional observational study methods.

Large electronic databases of medical records, billing codes, insurance claims, and prescriptions have enabled studies with larger sample sizes and a wider diversity of phenotypes under consideration.^4,5^ However, these databases are often restricted to billable events and may not capture phenotypes related to sub-clinical symptoms, behaviour, lifestyle, morphology, or over-the-counter medication.

Here, we sought to describe the broad spectrum of phenotypes associated with PD — the PD phenome — using a database that included 13,546 PD cases, >1·3 million controls, and 840 phenotypes including diagnoses, family history, medication usage, environmental exposures, and behaviours.

## Methods

### Study participants

People with PD were recruited through a targeted email campaign in conjunction with the Michael J. Fox Foundation and many other patient groups and clinics. In addition, individuals with PD were identified via online surveys from customers of the personal genetics company, 23andMe, Inc., who had consented to participate in research. All individuals provided informed consent, and 23andMe’s human subjects protocol was reviewed and approved by an AAHRPP-accredited institutional review board. This study was conducted according to the principles set out in the Declaration of Helsinki.

PD cases were individuals who self-reported having been diagnosed with PD. We excluded people if they reported a change in diagnosis or uncertainty about their diagnosis. We have previously shown that self-reported PD case status is accurate with 50 out of 50 cases confirmed via telemedicine interview.^6^ Age- and sex-matched controls were drawn from 23andMe research participants who self-reported that they have not been diagnosed with PD. For both PD cases and controls, we included unrelated individuals who were genetically inferred to have >97% European ancestry (see Supplementary Methods). We did this because 88·9% of all people with PD in the 23andMe database had >97% European ancestry and because PD incidence and prevalence may differ by ethnicity.^7,8^ We also removed individuals who self-reported having ever been diagnosed with: 1) an atypical parkinsonism or a non-parkinsonian tremor disorder; or 2) stroke, deep vein thrombosis, or pulmonary embolism (to reduce the probability of including individuals with vascular parkinsonism).

### Phenotypic data

Phenotypic data (including PD status) were collected using web or mobile-based surveys. Surveys were either based on existing instruments published in the medical literature, such as the “Reading the Mind in the Eyes” test^9^, or developed by 23andMe scientists, often in collaboration with medical professionals or a similarly relevant professional for non-medical surveys. Phenotype definitions are provided for phenotypes that were significantly associated with PD (Supplementary Table 1). Analyses were run on phenotypic data collected between November 10^th^, 2007 and October 10^th^, 2017.

**Table 1:** Demographics of PD cases and controls. The mean and standard deviation (SD) are presented for continuous phenotypes, while binary phenotypes are presented as a percentage. Sample sizes (N) vary between phenotypes due to differences in completion rates of various surveys. *P* values were not corrected for any other variables.

|  | Cases |  | Controls |  | *P* |
| --- | --- | --- | --- | --- | --- |
| | Mean (± SD) | N | Mean (± SD) | N | |
| Age | 69.8 (11.21) | 12,913 | 69.3 (11.39) | 116,217 | $7.29 \times 10^{-6}$ |
| Education years | 16.4 (2.77) | 7,890 | 16.3 (2.80) | 118,350 | $1.93 \times 10^{-3}$ |
| Body mass index | 26.5 (5.12) | 10,312 | 27.5 (5.19) | 144,368 | $8.78 \times 10^{-74}$ |
| Household income | \$118,571 (\$91,698) | 4,488 | \$136,483 (\$95,738) | 40,392 | $8.20 \times 10^{-33}$ |
| Female | 38.9% | 12,913 | 38.9% | 116,217 | 1 |
| Military service | 23.5% | 2,635 | 24.8% | 23,715 | 0.127 |
| Has smoked >100 cigarettes | 38.5% | 10,371 | 46.7% | 134,823 | $3.79 \times 10^{-58}$ |

### Covariates

All regressions were run using a standard set of covariates: age, sex, date of entry into the research cohort (split into 20 evenly populated bins, used to correct for differences in demographics and available surveys over time), and the first five genetic principal components (used to correct for differences in ancestry).

### Identifying significant, independent associations

For each of the 840 phenotypes, we selected individuals with non-missing data for the phenotype, covariates, and PD status. We then selected as many age-and sex-matched controls for each case as possible (see Supplementary Methods). Finally, we performed logistic regression with PD as the dependent variable and the phenotype in question as the independent variable, correcting for our standard covariates. In order to focus on independent phenotypes, we computed the partial correlation coefficient between each possible pair of phenotypes, a value ranging between -1 (perfectly anticorrelated) to 1 (perfectly correlated). For each pair of phenotypes where the absolute value of the partial correlation coefficient was greater than 0·25, we removed the phenotype that was less strongly associated with PD based on *P* value. To mitigate the risk of spurious associations with PD arising through multiple comparisons, we imposed a stringent Bonferroni-based *P* value threshold of 1·12x10^-4^ (0.05 / 446 independent phenotypes).

### Classification of associations based on previous literature

For each phenotype, we conducted a literature review and/or consulted the 23andMe Parkinson’s Disease Scientific Advisory Board to determine whether an association had previously been reported. Each association was systematically assigned to one of the following groups: known, likely, unclear, and novel (Supplementary Methods).

### Associations with disease duration in people with PD

To determine whether each significant phenotype passing Bonferroni correction was associated with PD duration, we ran regressions in PD cases with the phenotype as the outcome variable and PD duration and our standard covariates as the input variables. We used logistic regression for case-control phenotypes and linear regression for continuous phenotypes.

### Data sharing

Model outputs for all 840 logistic regressions can be found in Supplementary Table 2.

**Table 2:**
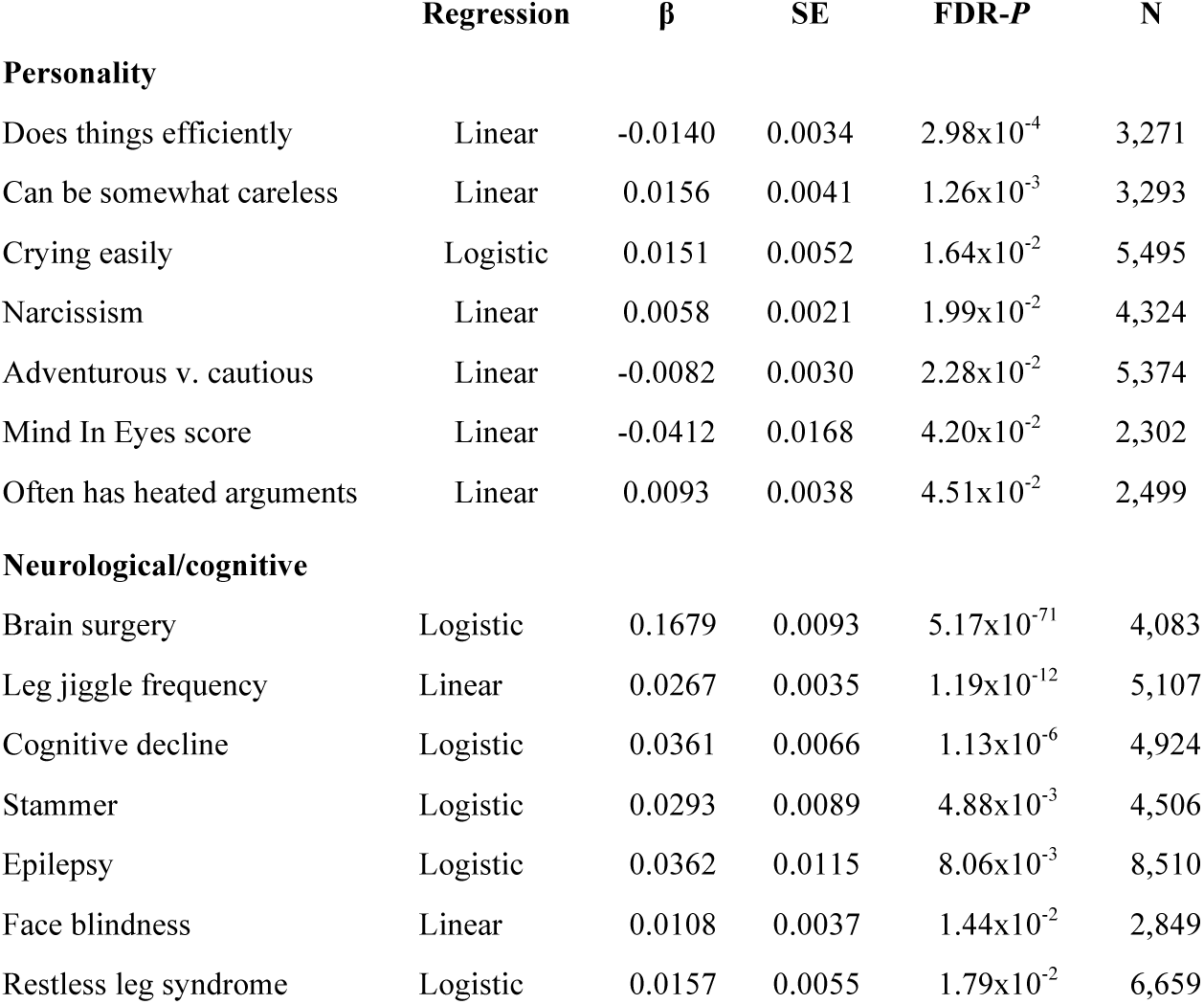
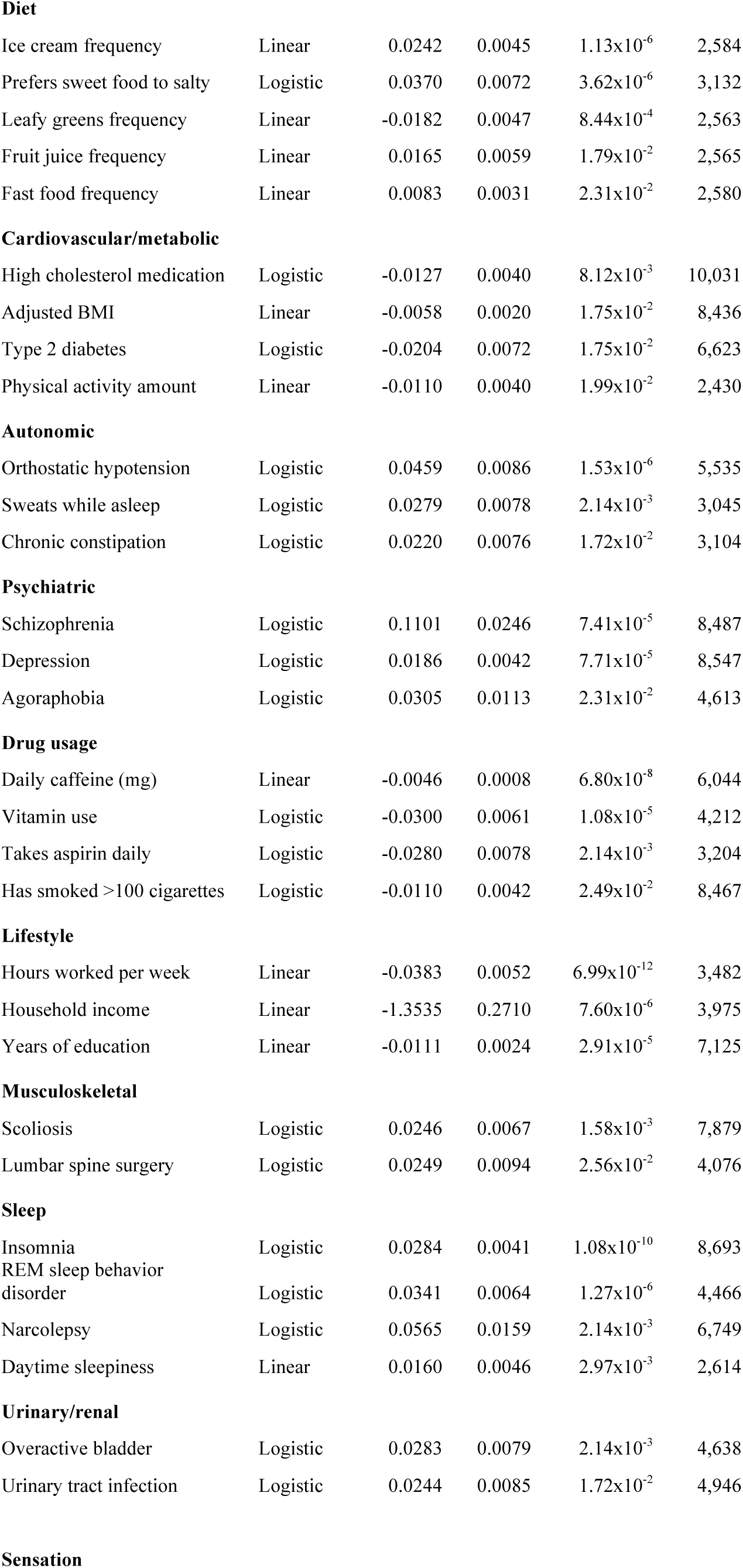
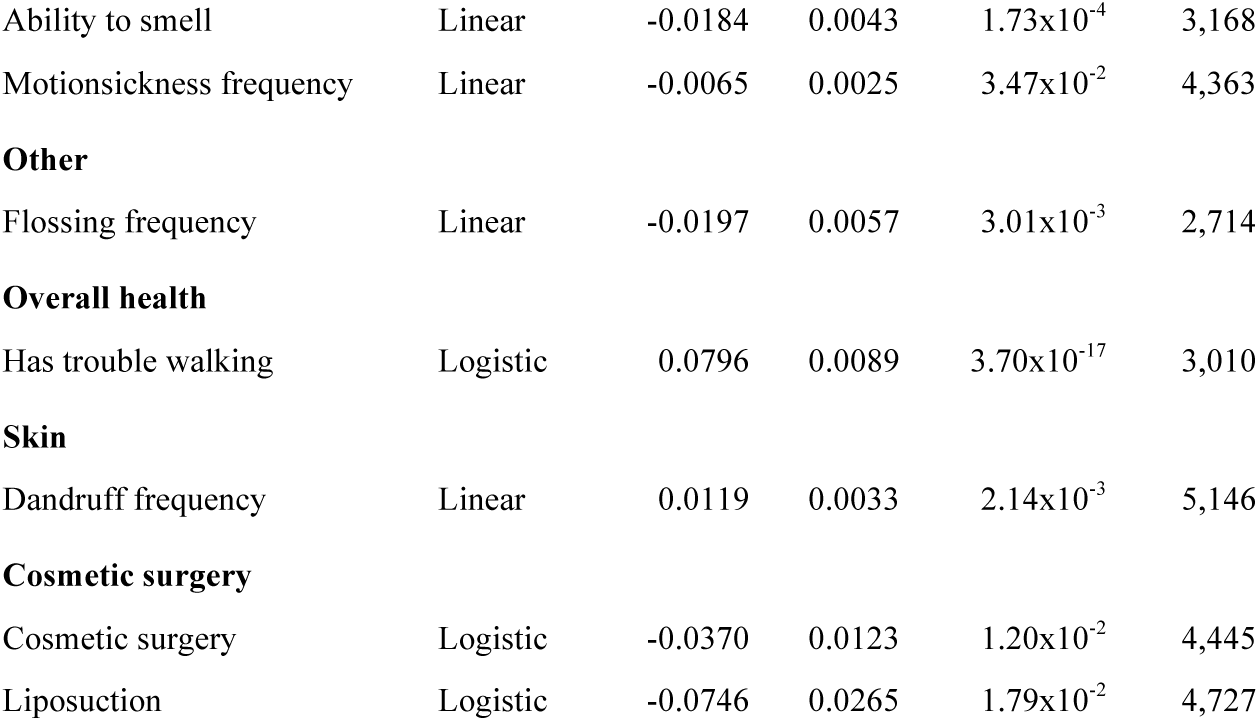
Phenotypes significantly associated with PD duration. We ran separate regressions for each of the 149 phenotypes that were significantly associated with PD case/control status using the phenotype as the dependent variable, PD duration as the independent variable, and correcting for our standard covariates. We used linear regressions for continuous phenotypes and logistic regressions for binary phenotypes. β = logarithm of the odds ratio for logistic regressions and effect estimate for linear regressions for a one year increase in PD duration, SE = standard error, FDR-*P* = false discovery rate-corrected *P* value, N = sample size.

| | Regression | $\beta$ | SE | FDR-*P* | N |
| --- | --- | --- | --- | --- | --- |
| **Personality** |  |  |  |  |  |
| Does things efficiently | Linear | -0.0140 | 0.0034 | $2.98 \times 10^{-4}$ | 3,271 |
| Can be somewhat careless | Linear | 0.0156 | 0.0041 | $1.26 \times 10^{-3}$ | 3,293 |
| Crying easily | Logistic | 0.0151 | 0.0052 | $1.64 \times 10^{-2}$ | 5,495 |
| Narcissism | Linear | 0.0058 | 0.0021 | $1.99 \times 10^{-2}$ | 4,324 |
| Adventurous v. cautious | Linear | -0.0082 | 0.0030 | $2.28 \times 10^{-2}$ | 5,374 |
| Mind In Eyes score | Linear | -0.0412 | 0.0168 | $4.20 \times 10^{-2}$ | 2,302 |
| Often has heated arguments | Linear | 0.0093 | 0.0038 | $4.51 \times 10^{-2}$ | 2,499 |
| **Neurological/cognitive** |  |  |  |  |  |
| Brain surgery | Logistic | 0.1679 | 0.0093 | $5.17 \times 10^{-71}$ | 4,083 |
| Leg jiggle frequency | Linear | 0.0267 | 0.0035 | $1.19 \times 10^{-12}$ | 5,107 |
| Cognitive decline | Logistic | 0.0361 | 0.0066 | $1.13 \times 10^{-6}$ | 4,924 |
| Stammer | Logistic | 0.0293 | 0.0089 | $4.88 \times 10^{-3}$ | 4,506 |
| Epilepsy | Logistic | 0.0362 | 0.0115 | $8.06 \times 10^{-3}$ | 8,510 |
| Face blindness | Linear | 0.0108 | 0.0037 | $1.44 \times 10^{-2}$ | 2,849 |
| Restless leg syndrome | Logistic | 0.0157 | 0.0055 | $1.79 \times 10^{-2}$ | 6,659 |
| **Diet** |  |  |  |  |  |
| Ice cream frequency | Linear | 0.0242 | 0.0045 | $1.13 \times 10^{-6}$ | 2,584 |
| Prefers sweet food to salty | Logistic | 0.0370 | 0.0072 | $3.62 \times 10^{-6}$ | 3,132 |
| Leafy greens frequency | Linear | -0.0182 | 0.0047 | $8.44 \times 10^{-4}$ | 2,563 |
| Fruit juice frequency | Linear | 0.0165 | 0.0059 | $1.79 \times 10^{-2}$ | 2,565 |
| Fast food frequency | Linear | 0.0083 | 0.0031 | $2.31 \times 10^{-2}$ | 2,580 |
| **Cardiovascular/metabolic** |  |  |  |  |  |
| High cholesterol medication | Logistic | -0.0127 | 0.0040 | $8.12 \times 10^{-3}$ | 10,031 |
| Adjusted BMI | Linear | -0.0058 | 0.0020 | $1.75 \times 10^{-2}$ | 8,436 |
| Type 2 diabetes | Logistic | -0.0204 | 0.0072 | $1.75 \times 10^{-2}$ | 6,623 |
| Physical activity amount | Linear | -0.0110 | 0.0040 | $1.99 \times 10^{-2}$ | 2,430 |
| **Autonomic** |  |  |  |  |  |
| Orthostatic hypotension | Logistic | 0.0459 | 0.0086 | $1.53 \times 10^{-6}$ | 5,535 |
| Sweats while asleep | Logistic | 0.0279 | 0.0078 | $2.14 \times 10^{-3}$ | 3,045 |
| Chronic constipation | Logistic | 0.0220 | 0.0076 | $1.72 \times 10^{-2}$ | 3,104 |
| **Psychiatric** |  |  |  |  |  |
| Schizophrenia | Logistic | 0.1101 | 0.0246 | $7.41 \times 10^{-5}$ | 8,487 |
| Depression | Logistic | 0.0186 | 0.0042 | $7.71 \times 10^{-5}$ | 8,547 |
| Agoraphobia | Logistic | 0.0305 | 0.0113 | $2.31 \times 10^{-2}$ | 4,613 |
| **Drug usage** |  |  |  |  |  |
| Daily caffeine (mg) | Linear | -0.0046 | 0.0008 | $6.80 \times 10^{-8}$ | 6,044 |
| Vitamin use | Logistic | -0.0300 | 0.0061 | $1.08 \times 10^{-5}$ | 4,212 |
| Takes aspirin daily | Logistic | -0.0280 | 0.0078 | $2.14 \times 10^{-3}$ | 3,204 |
| Has smoked >100 cigarettes | Logistic | -0.0110 | 0.0042 | $2.49 \times 10^{-2}$ | 8,467 |
| **Lifestyle** |  |  |  |  |  |
| Hours worked per week | Linear | -0.0383 | 0.0052 | $6.99 \times 10^{-12}$ | 3,482 |
| Household income | Linear | -1.3535 | 0.2710 | $7.60 \times 10^{-6}$ | 3,975 |
| Years of education | Linear | -0.0111 | 0.0024 | $2.91 \times 10^{-5}$ | 7,125 |
| **Musculoskeletal** |  |  |  |  |  |
| Scoliosis | Logistic | 0.0246 | 0.0067 | $1.58 \times 10^{-3}$ | 7,879 |
| Lumbar spine surgery | Logistic | 0.0249 | 0.0094 | $2.56 \times 10^{-2}$ | 4,076 |
| **Sleep** |  |  |  |  |  |
| Insomnia | Logistic | 0.0284 | 0.0041 | $1.08 \times 10^{-10}$ | 8,693 |
| REM sleep behavior disorder | Logistic | 0.0341 | 0.0064 | $1.27 \times 10^{-6}$ | 4,466 |
| Narcolepsy | Logistic | 0.0565 | 0.0159 | $2.14 \times 10^{-3}$ | 6,749 |
| Daytime sleepiness | Linear | 0.0160 | 0.0046 | $2.97 \times 10^{-3}$ | 2,614 |
| **Urinary/renal** |  |  |  |  |  |
| Overactive bladder | Logistic | 0.0283 | 0.0079 | $2.14 \times 10^{-3}$ | 4,638 |
| Urinary tract infection | Logistic | 0.0244 | 0.0085 | $1.72 \times 10^{-2}$ | 4,946 |
| **Sensation** |  |  |  |  |  |
| Ability to smell | Linear | -0.0184 | 0.0043 | $1.73 \times 10^{-4}$ | 3,168 |
| Motionsickness frequency | Linear | -0.0065 | 0.0025 | $3.47 \times 10^{-2}$ | 4,363 |
| **Other** |  |  |  |  |  |
| Flossing frequency | Linear | -0.0197 | 0.0057 | $3.01 \times 10^{-3}$ | 2,714 |
| **Overall health** |  |  |  |  |  |
| Has trouble walking | Logistic | 0.0796 | 0.0089 | $3.70 \times 10^{-17}$ | 3,010 |
| **Skin** |  |  |  |  |  |
| Dandruff frequency | Linear | 0.0119 | 0.0033 | $2.14 \times 10^{-3}$ | 5,146 |
| **Cosmetic surgery** |  |  |  |  |  |
| Cosmetic surgery | Logistic | -0.0370 | 0.0123 | $1.20 \times 10^{-2}$ | 4,445 |
| Liposuction | Logistic | -0.0746 | 0.0265 | $1.79 \times 10^{-2}$ | 4,727 |

## Results

Table 1 shows descriptive statistics for the demographics of PD cases and controls without adjusting for confounding variables. Because we used age-and sex-matched controls, PD cases and controls were similar with respect to average age (cases: 69·8 ± 11·2 SD, controls: 69·3 ± 11·4 SD) and sex ratio (cases: 38·9% female, controls: 38·9% female). We found no significant difference between PD cases and controls for education (*P* = 1·94x10^-3^) or military service (*P* = 1·27x10^-1^) after correcting for multiple testing. Compared to controls, people with PD had a lower body mass index (cases: 26·5 ± 5·1 SD, controls: 27·5 ± 5·2 SD, *P* = 8·78x10^-74^), lower household income (cases: $118,571 ± $91,698 SD, controls: $136,483 ± $95,738 SD, *P* = 8·20x10^-33^), and were less likely to have ever smoked more than 100 cigarettes (cases: 38·5%, controls: 46·7%, *P* = 3·79x10^-58^). The percentage of cases and controls living in various United States Census Bureau divisions was similar (Supplementary Figure 1).

**Figure 1.**
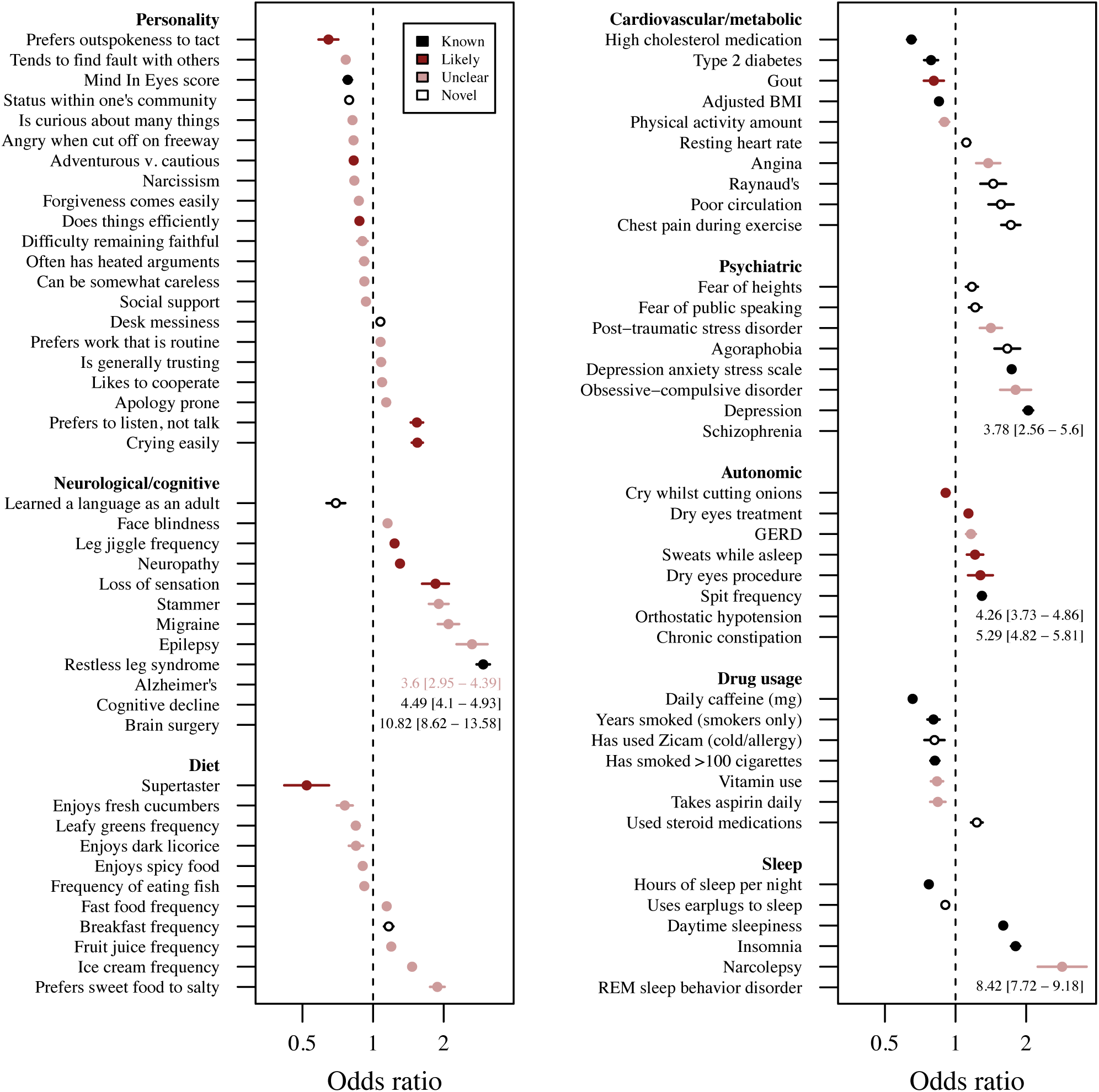
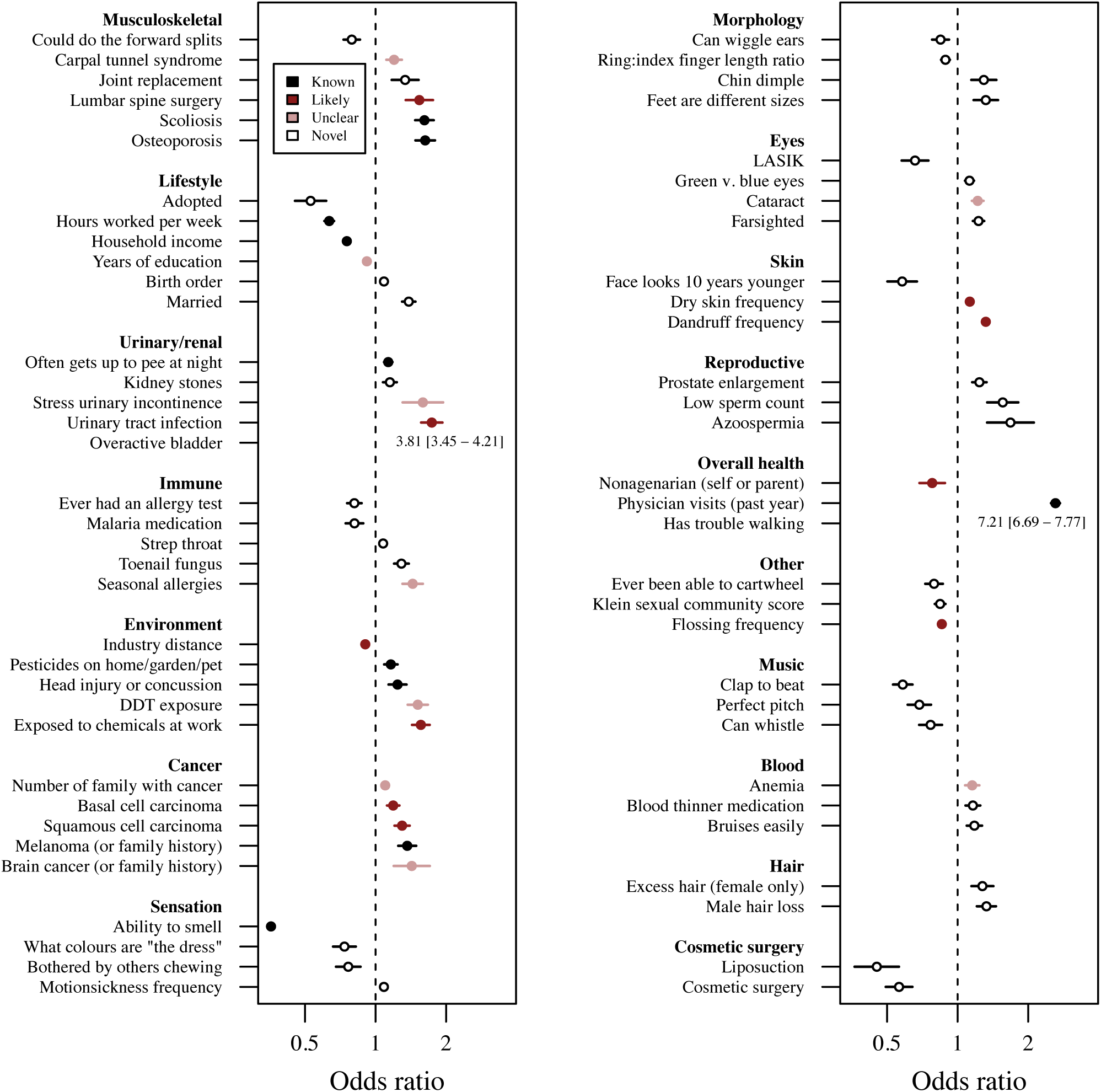
Forest plot of the odds ratios (ORs) ± 95% confidence intervals (CIs) for 149 phenotypes that were significantly associated with PD, sorted by group and OR. Colours denote the extent to which an association between PD and the phenotype had previously been studied. If the upper 95% CI of an association was greater than four, data are presented as text. The dashed line represents an OR of one—no association with PD. To help clarify the directionality of associations, comprehensive phenotype definitions can be found in Supplementary Table 1. BMI = body mass index, GERD = gastro-esophageal reflux disease, mg = milligrams, REM = rapid eye movement, DDT = dichlorodiphenyltrichloroethane, LASIK = laser-assisted in situ keratomileusis.

After Bonferroni-correction (*P* < 1·12x10^-4^) there were 149 independent phenotypes that were significantly associated with PD (Figure 1). We were able to replicate 32 “known” associations (see Supplementary Methods) that have previously been reported in the literature, demonstrating that our online self-reported cohort is phenotypically similar to previously described clinical cohorts. For example, we replicated the negative association between caffeine intake and PD (OR = 0·657 [95% CI 0·635 – 0·679], *P* = 3·01x10^-139^), and the positive association between PD and constipation (OR = 5·29 [95% CI 4·82 – 5·81], *P* = 8·45x10^-271^). The only known association that we failed to detect was a decreased prevalence of cancer (excluding skin cancer) in PD cases. The size of our cohort and the diversity of phenotypes studied also allowed us to discover 49 novel associations that, to our knowledge, have not been previously reported. We also found 24 phenotypes that were strongly related to known aspects of PD (“likely”) and 44 phenotypes where an association with PD has previously been studied, but did not meet our criteria for being classified as “known” or “likely” (“unclear”). Twenty-six phenotypes were significantly associated with disease duration in people with PD (Table 2, Supplementary Table 3).

## Discussion

Although Parkinson’s disease is diagnosed on the basis of motor signs, the PD phenome is known to encompass a broad range of non-motor features such as autonomic dysfunction, sleep disturbances, cognitive dysfunction, and psychiatric disorders. By systematically analysing a large database of phenotypic data, we replicated many of these established associations and discovered several novel associations. We tested 840 phenotypes for an association with PD — 446 of which were uncorrelated with each other — and found 149 significant associations after correcting for multiple testing. The odds ratios (ORs) for significant positive associations (OR > 1) ranged from 1.075 (“Desk messiness”) to 10.8 (“Brain surgery”), while significant negative associations (OR < 1) ranged from 0.933 (“Social support”) to 0.358 (“Ability to smell”).

It is important to note that phenotypic associations can arise due to a number of different underlying relationships including causation, reverse causation, and confounding (Supplementary Figure 2). Our goal for this study was to meaningfully extend the characterization of the PD phenome, not to make causal inferences from cross-sectional data. We have organized PD-associated traits into 25 subjective functionally related groups and discuss some of the findings here.

### Personality

People with PD tended to report being less argumentative, less quick to anger, less outspoken, and less talkative. It is difficult to compare personality traits between different studies, but our results are qualitatively similar to previous observations, which have found that people with PD tended to be less aggressive^10^, more inhibited^11^, and more neurotic and introverted.^12^ Whether these personality differences exist prior to PD diagnosis is controversial and may be due to recall bias.^13^ We found that 14 of the 21 personality phenotypes (67%) were not associated with PD duration (false discovery rate-corrected *P* [FDR-*P*] > 0·05). This suggests that many of these phenotypes do not arise in the later stages of PD, but rather may arise in early PD or before diagnosis. Many of the personality phenotypes that were significantly associated with PD duration are plausibly linked to more advanced PD, such as doing things less efficiently, being less narcissistic, and being more careless.

### Diet

People with PD were more likely to report a preference for, and increased consumption of, sweet foods and beverages. There have been few consistent associations between PD and the intake of specific food items with the exception of coffee^14^ and perhaps dairy products.^15^ However, a recent Italian study found a greater intake of sweets in 600 people with PD compared to 600 age- and sex-matched controls.^16^ A German case-control study including 342 people with PD reported a greater intake of sweets, cookies, and cakes^17^ and that this was associated with time since diagnosis. We also found PD duration was associated with increased ice cream and fruit juice consumption and an increased probability of preferring sweet foods over salty foods (FDR-*P* < 0·05). These results suggest that altered food preferences may be a feature of later stage PD.

Despite a greater consumption of sweet foods, people with PD were less likely to have diabetes or take diabetes medication. This observation is consistent with several case-control studies that also found a negative relationship between PD and diabetes.^14^ However, longitudinal cohort studies have mainly found positive associations.^14^ The reason for this discrepancy is unclear and is important to clarify through further study.

### Neurological and cognitive phenotypes

Several neurological disorders were associated with PD. Restless leg syndrome (RLS) is more prevalent in people with PD, although prevalence estimates range widely and it is unclear whether RLS is a risk factor for PD.^18^ The pathological hallmarks of Alzheimer’s disease are more common in individuals diagnosed with PD^19^, and cognitive decline is a feature of PD that may occur early or late in the disease.^1^

We observed that migraine was positively associated with PD. Only two cohort studies have previously been conducted, but both found that midlife migraine was associated with increased risk of PD.^20,21^ However, these studies have been criticized for not correcting for cardiovascular disease, head injury, and/or other neurological diseases. We found that the association between PD and migraine remained nominally significant (OR = 1·205 [1·056 – 1·374], *P* = 5·49x10^-3^) after correcting for possible confounding factors (see Supplementary Methods). Since the average age of onset was typically much earlier for migraine than for PD, migraine may be a novel risk factor for PD. Indeed, we found that migraine was not significantly associated with PD duration (FDR-*P* = 0·832).

People with PD were more likely to self-report having been diagnosed with, or treated for, epilepsy or seizures. A recent British study found a significantly greater incidence rate ratio of epilepsy in 23,086 PD cases and 92,343 matched controls.^22^ Similarly, we found that these self-reported seizures were significantly associated with PD duration (FDR-*P* = 8·06x10^-3^), suggesting they may arise later in the disease process.

### Psychiatric disorders

Psychiatric manifestations are common in PD. Questions pertaining to depression and anxiety have been incorporated into the Movement Disorder Society (MDS)-sponsored revision of the Unified Parkinson’s Disease Rating Scale (UPDRS)^1^ and psychosis has been found in up to 60% of PD patients.^23^ There has been substantial interest in a connection between obsessive-compulsive disorder (OCD) and PD since both can be treated via deep brain stimulation, can be caused by viral encephalitis, and may involve dysfunction of the basal ganglia.^24^ Kummer and Teixeira^24^ reviewed 11 studies of OCD and OCD symptoms (OCS) in relation to PD and found several methodological issues including small sample sizes (20 – 124 PD cases) leading to a lack of statistical power. Indeed, four of the five case-control studies assessed yielded non-significant results. With 10,574 PD cases in this particular regression, our study is the largest analysis of PD and OCD comorbidity to date. We observed a positive association between PD and OCD (*P* = 2·00x10^-14^), and that OCD was not associated with PD duration (FDR-*P* = 0·182).

### Lifestyle factors

Being married was associated with an increased odds of PD. Compared to people who reported never having been married (Supplementary Figure 3) and after adjusting for age and our other covariates, the odds of PD were: 1) highest in people who were currently married (OR = 1·793 [1·609 – 1·999], *P* = 5·04x10^-26^), 2) attenuated in people who were divorced (OR = 1·268 [1·108 – 1·450], *P* = 5·40x10^-4^), and 3) abrogated in people who were widowed (OR = 1·029 [0·852 – 1·241], *P* = 0·771). We ran our phenotype association pipeline with marital status as the outcome variable and found that marriage was largely negatively associated with a wide variety of diseases, with PD being a notable exception (Supplementary Table 4). This remained true when analyzing females and males separately. These results suggest that the association with PD was unlikely to be due to bias from over-reporting of disease history in married individuals, but we cannot completely rule out the possibility that prodromal features of PD like REM sleep behaviour disorder may lead spouses to encourage their partners to seek medical consultation. Marital status was not significantly associated with PD duration (FDR-*P* = 0·743).

### Other comorbidities

We found that seasonal allergies were associated with PD. Although many allergies were significantly correlated with one another, only a subset of these allergies was associated with PD. PD was associated with allergies to plants and antibiotics, but not with allergies to food or animals (Supplementary Table 2). To our knowledge, only two studies of PD and allergies have been published. A retrospective case-control study found that PD was positively associated with allergic rhinitis, but not asthma or hayfever.^25^ A study using the Taiwan National Health Insurance Research Database found a positive, but not significant, association between PD and allergic rhinitis (hazard ratio = 1·29 [0·97–1·72]). Although little research has been done on PD and allergies, the immune system is known to play an important role in PD. For example, a recent meta-analysis of 25 case-control studies found that the peripheral concentration of several inflammatory cytokines was increased in people with PD.^26^ Seasonal allergies typically develop early in life, but further research is needed to determine whether having seasonal allergies is a risk factor for PD. If so, it may be possible to manipulate by risk using immune-modulating drugs. Indeed, we found that allergies were not significantly associated with PD duration (FDR-*P* = 0·252).

We found a significant positive association between PD and anemia. A Mendelian randomization study using 23andMe data found evidence that decreased serum iron levels are causally associated with increased PD risk.^27^ A study of incident PD cases from the Rochester Epidemiology Project found that anemia was associated with PD, particularly when the anemia diagnosis preceded the PD diagnosis by 20-29 years. Furthermore, we found that anemia was not associated with PD duration (FDR-*P* = 0·356). These results suggest that serum iron levels may be a modifiable risk factor for PD. It should be noted that a meta-analysis of ten studies^28^ found that serum iron levels were not significantly different in people with PD; however, there was significant heterogeneity between studies (I^2^ = 0·93, *P* = 0·0001).

### Limitations

There were several limitations to our study. First, PD diagnoses were self-reported and not confirmed by a clinician. However, we have previously shown that a telemedicine-based assessment by a neurologist confirmed self-reported PD status in 100% of a sample of fifty people with PD from the 23andMe cohort.^6^ Second, our analyses were restricted to individuals of European ancestry and further work is needed to determine whether our results are applicable to individuals from other ethnic backgrounds. We acknowledge that there is a diversity problem in medical research and we have begun several initiatives to address this in our research. Third, although our regressions were controlled for age, sex, ancestry, and date of enrollment into the 23andMe research cohort, there may be other confounding variables that we have not included. For example, people with PD were given 23andMe kits for free while controls were drawn from paying customers. We did not correct for the difference in socio-economic status between PD cases and controls because we only had household income data for 35% of PD cases, but the effect estimates from significant associations were strongly correlated with the corresponding effect estimates from models that included household income as a covariate (*r* = 0·985, *P* = 4·7x10^-87^). Finally, survey completion rates were variable across phenotypes. Because we discarded individuals with missing phenotypic data, each regression was run on a different subset of individuals. This makes it more difficult to directly compare the odds ratios for different phenotypes. However, all regressions were run using consistent methodology and are more comparable than values derived from completely separate studies.

## Summary

We characterized the PD phenome using a systematic screen of a large database of online survey responses. We replicated 32 known associations, demonstrating that our self-reported PD cases were phenotypically similar to PD cases from other studies. We also discovered 49 associations that have not previously been reported, which will need to be replicated in a separate data set. Many personality traits were associated with PD, but the mechanisms linking PD and personality are unknown. Several phenotypes related to an increased preference for or consumption of sweet foods and beverages were associated with PD duration and may therefore be consequences of PD progression. We found that migraine, seasonal allergies, and anemia were associated with PD, all of which tend to have an age of onset several decades before the average age of diagnosis for PD. More research is needed to determine precisely which aspects of the PD phenome exist earlier in life. Early features of PD may be used to improve PD prediction algorithms and may represent modifiable PD risk factors that could be targeted therapeutically.

### Contributors

KH, PC, PF, and BA designed the study. KH performed all statistical analyses. KH and AJN conducted the literature review. Members of the 23andMe Research Team provided support and infrastructure to enable data collection and the analyses presented here. All authors assisted with analyzing and interpreting study results. KH wrote the first draft of the manuscript. All authors assisted with editing subsequent drafts.

### 23andMe Research Team

Michelle Agee, Adam Auton, Robert K. Bell, Katarzyna Bryc, Sarah L. Elson, Nicholas A. Furlotte, David A. Hinds, Jennifer C. McCreight, Karen E. Huber, Aaron Kleinman, Nadia K. Litterman, Matthew H. McIntyre, Joanna L. Mountain, Elizabeth S. Noblin, Carrie A.M. Northover, Steven J. Pitts, J. Fah Sathirapongsasuti, Olga V. Sazonova, Janie F. Shelton, Suyash Shringarpure, Chao Tian, Joyce Y. Tung, Vladimir Vacic, Catherine H. Wilson

## Acknowledgements

We thank the customers of 23andMe who answered surveys and participated in this research. We thank the 23andMe Parkinson’s Disease Scientific Advisory Board — Alice Chen-Plotkin, David Standaert, Owen Ross, Ray Dorsey, and Uta Francke — for their helpful comments during the preparation of the manuscript

## Tables

**Supplementary Table 1.** Phenotype definitions for 149 independent phenotypes that were significantly associated with PD, sorted by *P* value from lowest to highest. For each we provide an internal name, publication name, and definition. The internal name is the identifier used to tag each phenotype in the 23andMe Research Environment. The publication name is the identifier used to represent each phenotype throughout this study and is intended to be short and intuitive. The definition section describes how each phenotype was defined. Where relevant, we have included all or a subset of possible responses in parentheses at the end of the definition. If there were three or fewer possible responses, all are listed separated by commas. If there were greater than three possible responses (typically five), the most extreme options are presented separated by a hyphen.

**Supplementary Table 2.** Logistic regression model outputs for all 850 phenotypes. Results are organized into 446 groups of correlated phenotypes, each separated by an empty row. Sample size is reported separately for PD cases (“n_pd”) and controls (“n_controls”). Two prevalences are reported for binary phenotypes: “prevalence_total”, the prevalence of the phenotype in the entire 23andMe database, and “prevalence_merged”, the prevalence of the phenotype in the 23andMe database after removing individuals missing data for PD or covariates. Standard deviation is presented for continuous phenotypes.

**Supplementary Table 3.** Associations between PD duration and 149 phenotypes that were significantly associated with PD case-control status. Test type = whether linear or logistic regression was used. Beta = log odds ratio for logistic regression; effect size for linear regression. FDR-corrected *P* value = false discovery rate-corrected *P* value. N = sample size

**Supplementary Table 4.** Diseases that were negatively (A) or positively (B) associated with being married. We used the same phenotype regression pipeline as for the Parkinson’s disease analysis, except we did not use age-and sex-matched controls. Instead, we included five additional age covariates, age raised to the power of two to age raised to the power of six. We excluded phenotypes that: 1) had a *P* value greater than that for PD and marriage (5.85x10^-42^), 2) were not diseases, 3) were infections or injuries, and 4) were quantitative (not case-control). There were many more diseases that were negatively associated (24 diseases) than were positively associated (6 diseases) with marriage. Many of the negatively associated diseases were psychiatric (9) or neurological (4). Two of the diseases that were positively associated with marriage were related to infertility, which is probably due to reporting bias of married couples being more likely to try to get pregnant. There were also two positively-associated diseases related to skin cancer. n = sample size. prevalence_total = the prevalence of the phenotype in the entire 23andMe database. prevalence_merged = the prevalence of the phenotype in the 23andMe database after removing individuals missing data for marital status and covariates. t2d_broad = Type 2 diabetes, iqb.ivf = in vitro fertilization.

## Figure Legends

**Supplementary Figure 1.** The proportion of PD cases (light grey) and controls (dark grey) living in each of the nine United States Census Bureau divisions.

**Supplementary Figure 2.** Directed acyclic graphs displaying several different scenarios that can lead to an association between PD and another phenotype. Black solid arrows denote a causal relationship and grey dashed arrows denote a spurious association. The other phenotype could cause PD (A), PD could cause the other phenotype (B), or the association could be driven by a third confounding variable that is associated with both PD and the other phenotype (C). These scenarios are not necessarily mutually exclusive. It is important to remember that, even when a study has attempted to take potentially confounding variables into account, there may be other unmeasured confounders that have not been considered. The aim of this study was to identify phenotypes that were associated with PD regardless of the underlying relationship that led to the association. In future, we plan to assess causality for these phenotypes using various epidemiological tools, including Mendelian randomization.

**Supplementary Figure 3.** The odds ratio (± 95% confidence interval) of having PD for various marital statuses as compared to individuals who have never been married.

## Supplementary Methods

### Study participants

Individuals with PD were recruited through a targeted email campaign in conjunction with the Michael J. Fox Foundation, The Parkinson’s Institute and Clinical Center, and many other PD patient groups and clinics. Emails or hard copy mailings were sent to all individuals who had registered with these groups as PD patients. For a fuller description of recruitment, screening, and genotyping see Do *et al*.^1^

### Institutional review board

All individuals answered surveys online according to 23andMe’s human subjects protocol, which was reviewed and approved by Ethical & Independent Review Services, an AAHRPP-accredited institutional review board.

### Determining unrelated individuals of European ancestry

Individuals were only included if they had >97% European ancestry, as determined through an analysis of local ancestry.^2^ Briefly, this analysis first partitions phased genomic data into short windows of ∼100 SNPs. Within each window, a support vector machine is used to classify individual haplotypes into one of 31 reference populations. The support vector machine classifications are then fed into a hidden Markov model (HMM) that accounts for switch errors and incorrect assignments, and gives probabilities for each reference population in each window. Finally, simulated admixed individuals are used to recalibrate the HMM probabilities so that the reported assignments are consistent with the simulated admixture proportions. The reference population data are derived from public data sets (the Human Genome Diversity Project, HapMap and 1000 Genomes) and from 23andMe research participants who have reported having four grandparents from the same country.

A maximal set of unrelated individuals was chosen for the analysis using a segmental identity-by-descent estimation algorithm^3^. Individuals were defined as related if they shared 4700 cM identity-by-descent, including regions where the two individuals share either one or both genomic segments identical-by-descent. This level of relatedness (roughly 20% of the genome) corresponds approximately to the minimal expected sharing between first cousins in an outbred population.

### Selection of phenotypes

We conducted a systematic analysis of a curated set of 840 phenotypes in the 23andMe database. All phenotypes for which we have conducted a genome-wide association study since January 2016 were included, as were all phenotypes that are considered sufficiently important to warrant storage of monthly data snapshots. We excluded phenotypes pertaining to drug side effects, covariates used in downstream analyses, and phenotypes for which PD was an inclusion or exclusion criterion. Survey completion rates varied across phenotypes. After removing individuals with missing data for PD case-control status, covariates, and the phenotype in question, we excluded phenotypes for which there were 1) fewer than 2,000 PD cases, 2) fewer than 10,000 PD cases or controls, or 3) fewer than 30 individuals with both PD and the phenotype in question. Finally, we excluded phenotypes where we could not assign at least one age-and sex-matched control to each PD case.

### Age-and sex-matched controls

We 1) separated cases by sex and divided each into ten age bins; 2) determined the number of controls that fell into each corresponding sex/age bin; 3) divided the number of controls in a bin by the number of corresponding cases; 4) determined *n*, the minimum number of available controls per case across all bins; and 5) randomly selected *n* age-and sex-matched controls for each case.

### Calculating pairwise partial correlation coefficients

All 23andMe phenotypes are internally labeled with one or more “tags” in order to group related phenotypes (*e.g.* neurological). In order to make the calculation of pairwise partial correlation coefficients more computationally efficient, we only computed values for pairs of phenotypes with the same tag. At the end, we computed pairwise partial correlation coefficients for all significant phenotypes to remove correlated phenotypes that did not have the same tag.

### Correcting for multiple comparisons

We used a conservative Bonferroni threshold to determine statistical significance for our phenotypic association tests in order to minimize the risk of Type I error. Since we studied 446 independent phenotypes, we used a *P* value threshold of 0.05 / 446 = 1·12x10^-4^. For phenotypes that were significantly associated with PD, we had a strong *a priori* reason to expect that they may also be associated with PD duration. Therefore, we used a less conservative false discovery rate (FDR) threshold to determine significance for our regressions with PD duration. Specifically, we used the p.adjust function in R using the “FDR” method to obtain FDR-corrected *P* values.

### Classification of associations based on previous literature

We used PubMed to perform a MEDLINE database search using the keyword “Parkinson” and various keywords to describe each significant phenotype. For example, for the phenotype “blood thinner medication” we performed separate searches using the following keywords: “blood thinner”, “anticoagulant”, “warfarin”, and “platelet”. We supplemented these PubMed searches with simple web searches as well. We identified meta-analyses, review articles, cohort studies, and case-control studies that tested for an association between the prevalence (or incidence) of PD and the prevalence (or incidence) of the phenotype in question. We extracted information regarding these associations from publications that met our search criteria. For each association, one of the authors (KH) manually assessed the results from all relevant publications and assigned the association to one of four categories denoting the extent to which the association has previously been studied: known, likely, unclear, and novel. Another author (AJN) reviewed these assignments and discrepancies in assignments between the authors were discussed until consensus was reached. The 23andMe PD Scientific Advisory Board assessed a subset of associations.

“Known” associations were those that have repeatedly and consistently been found to be associated with PD or those that are well-known in the clinical community. Specifically, an association was classified as “known” if: 1) the phenotype is used in the Movement Disorder Society criteria for PD^4^ or prodromal PD^5^, or the MDS-UPDRS^6^; 2) a meta-analysis has found a significant association; 3) a review article by domain experts asserted that an association is known; or 4) the association is caused by a known difference in how PD cases and controls were recruited. “Likely” associations were those where we were unable to find previous literature that demonstrated an association, but where we achieved consensus agreement that the phenotype was strongly related to “known” aspects of PD. For example, we were unable to find any literature showing that, on average, tooth flossing frequency is decreased in PD patients, but this is likely given the motor complications of PD. “Unclear” associations were those that have been previously been reported in the literature, but for which no clear consensus exists. This could be because: 1) different studies found evidence of an association with opposite directions of effect, 2) the relevant studies were underpowered, or 3) concerns have been raised over the validity of the association due to potentially confounding variables. “Novel” associations were ones for which we could not find any relevant literature.

### Migraine: correcting for potential confounders

Previous studies have found an association between PD and migraine, but have been criticized for not correcting for potential confounding variables such as head injury, other neurological diseases, and cardiovascular risk factors and outcomes. We re-ran the regression for migraine after including phenotypes related to neurological disease, head injury, and cardiovascular disease that were associated with both PD and migraine with the same direction of effect: angina, chest pain during exercise, blood thinner use, neuropathy, epilepsy, restless leg syndrome, and REM sleep behaviour disorder. In addition, we included several other potentially confounding phenotypes that were significantly associated with both PD and migraine with the same direction of effect: depression, insomnia, anaemia, constipation, and the Depression Anxiety Stress Scale score. To account for missing data, we created dummy variables for each phenotype that encoded individuals with missing data as one and non-missing individuals as zero.

## References

1. Goetz CG, Tilley BC, Shaftman SR, et al. Movement Disorder Society-Sponsored Revision of the Unified Parkinson’s Disease Rating Scale (MDS-UPDRS): Scale presentation and clinimetric testing results. Mov Disord 2008; 23: 2129–70.

2. Wirdefeldt K, Adami H-O, Cole P, et al. Epidemiology and etiology of Parkinson’s disease: a review of the evidence. Eur J Epidemiol 2011; 26: 1–58.

3. Hirtz D, Thurman D, Gwinn-Hardy K, Mohamed M, Chaudhuri A, Zalutsky R. How common are the ‘common’ neurologic disorders? Neurology 2007; 68: 326–37.

4. Frandsen R, Kjellberg J, Ibsen R, Jennum P. Morbidity in early Parkinson’s disease and prior to diagnosis. Brain Behav 2014; 4: 446–52.

5. Nielsen SS, Warden MN, Wright BA, Racette BA. A predictive model to identify Parkinson disease from administrative claims data. Neurology 2017; 89: 1448–56.

6. Dorsey ER, Darwin KC, Mohammed S, et al. Virtual research visits and direct-to-consumer genetic testing in Parkinson’s disease. Digit Heal 2015; 0: 1–18.

7. Van Den Eeden SK, Tanner CM, Bernstein AL, et al. Incidence of Parkinson’s Disease: Variation by Age, Gender, and Race/Ethnicity. Am J Epidemiol 2003; 157: 1015–1022.

8. Wright Willis A, Evanoff BA, Lian M, Criswell SR, Racette BA. Geographic and ethnic variation in Parkinson disease: A population-based study of us medicare beneficiaries. Neuroepidemiology 2010; 34: 143–51.

9. Baron-Cohen S, Wheelwright S, Hill J, Raste Y, Plumb I. The Reading the Mind in the Eyes Test Revised Version?: A Study with Normal Adults, and Adults with Asperger Syndrome or High-functioning Autism. J Child Psychol Psychiat Assoc Child Psychol Psychiatry 2001; 42: 241–51.

10. Todes CJ, Lees AJ. The pre-morbid personality of patients with Parkinson’s disease. J Neurol Neurosurg Psychiatry 1985; 48: 97–100.

11. Heberlein I, Ludin HP, Scholz J, Vieregge P. Personality, depression, and premorbid lifestyle in twin pairs discordant for Parkinson’s disease. J Neurol Neurosurg Psychiatry 1998; 64: 262–6.

12. Sieurin J, Gustavsson P, Weibull CE, et al. Personality traits and the risk for Parkinson disease: a prospective study. Eur J Epidemiol; 31: 169–175.

13. Poletti M, Bonuccelli U. Personality traits in patients with Parkinson’s disease: Assessment and clinical implications. J Neurol 2012; 259: 1029–38.

14. Noyce AJ, Bestwick JP, Silveira-Moriyama L, et al. Meta-analysis of early nonmotor features and risk factors for Parkinson disease. Ann Neurol 2012; 72: 893–901.

15. Chen H, O’Reilly E, McCullough ML, et al. Consumption of dairy products and risk of parkinson’s disease. Am J Epidemiol 2007. DOI:10.1093/aje/kwk089.

16. Cassani E, Barichella M, Ferri V, et al. Dietary habits in Parkinson’s disease: Adherence to Mediterranean diet. Parkinsonism Relat Disord 2017; 42: 40–6.

17. Hellenbrand W, Seidler A, Boeing H, et al. Diet and Parkinson’s disease. I: A possible role for the past intake of specific foods and food groups. Results from a self-administered food-frequency questionnaire in a case-control study. Neurology 1996; 47: 636–43.

18. Högl B, Stefani A. Restless legs syndrome and periodic leg movements in patients with movement disorders: Specific considerations. Mov Disord 2017; 32: 669–81.

19. Hughes AJ, Daniel SE, Kilford L, Lees AJ. Accuracy of clinical diagnosis of idiopathic Parkinson’s disease: a clinico-pathological study of 100 cases. J Neurol Neurosurg Psychiatry 1992; 55: 181–4.

20. Scher AI, Ross GW, Sigurdsson S, et al. Midlife migraine and late-life parkinsonism: AGES-reykjavik study. Neurology 2014; 83: 1246–1252.

21. Wang H-I, Ho Y-C, Huang Y-P, Pan S-L. Migraine is related to an increased risk of Parkinson’s disease: A population-based, propensity score-matched, longitudinal follow-up study. Cephalalgia 2016; 36: 1316–23.

22. Gruntz K, Bloechliger M, Becker C, et al. Parkinson’s disease and the risk of epileptic seizures. Ann Neurol 2018; : 363–74.

23. Fénelon G, Soulas T, Zenasni F, De Langavant LC. The changing face of Parkinson’s disease- associated psychosis: A cross-sectional study based on the new NINDS-NIMH criteria. Mov Disord 2010; 25: 763–6.

24. Kummer A, Teixeira AL. Parkinson’s Disease and Obsessive-Compulsive Phenomena: A Systematic Review. Curr Psychiatry Rev 2009; 5: 55–61.

25. Bower J, Maraganore D, Peterson B, Ahlskog J, Rocca W. Immunologic diseases, anti-inflammatory drugs, and Parkinson disease: a case-control study. Neurology 2006; 67: 494–6.

26. Qin X-Y, Zhang S-P, Cao C, Loh YP, Cheng Y. Aberrations in Peripheral Inflammatory Cytokine Levels in Parkinson Disease. JAMA Neurol 2016; 73: 1316–24.

27. Pichler I, Del Greco MF, Gögele M, et al. Serum Iron Levels and the Risk of Parkinson Disease: A Mendelian Randomization Study. PLoS Med 2013; 10: e1001462.

28. Mariani S, Ventriglia M, Simonelli I, et al. Fe and Cu do not differ in Parkinson’s disease: A replication study plus meta-analysis. Neurobiol Aging 2013; 34: 632–3.

## References

1. Do CB, Tung JY, Dorfman E, et al. Web-based genome-wide association study identifies two novel loci and a substantial genetic component for Parkinson’s disease. PLoS Genet 2011; 7: e1002141.

2. Durand EY, Do CB, Mountain JL, Macpherson JM. Ancestry Composition: A Novel, Efficient Pipeline for Ancestry Deconvolution. bioRxiv 2014. doi:10.1101/010512.

3. Henn BM, Hon L, Macpherson JM, et al. Cryptic Distant Relatives Are Common in Both Isolated and Cosmopolitan Genetic Samples. 2012; 7. doi:10.1371/journal.pone.0034267.

4. Postuma RB, Berg D, Stern M, et al. MDS clinical diagnostic criteria for Parkinson’s disease. Mov Disord 2015; 30: 1591–601.

5. Berg D, Postuma RB, Adler CH, et al. MDS research criteria for prodromal Parkinson’s disease. Mov Disord 2015; 30: 1600–11.

6. Goetz CG, Tilley BC, Shaftman SR, et al. Movement Disorder Society-Sponsored Revision of the Unified Parkinson’s Disease Rating Scale (MDS-UPDRS): Scale presentation and clinimetric testing results. Mov Disord 2008; 23: 2129–70.

